# Dynamic Plk1 recruitment to the inner centromere

**DOI:** 10.1101/2024.07.03.601947

**Authors:** Roshan X Norman, Robert F Lera, Anuoluwapo A Mattix, Zhouyuan Shen, Caleb L Carlsen, Mark E Burkard

## Abstract

Mitosis is carefully orchestrated by reversible phosphorylation events. Polo-like kinase 1 (Plk1) regulates multiple functions across the kinetochore during mitotic progression. Recently, Bub1 (outer kinetochore) and CENP-U (inner kinetochore) were described as two major sites of Plk1 recruitment to the kinetochore. Here, we report an additional dynamic site of Plk1 recruitment to the inner centromere. Inner centromere docking occurs during late prometaphase and metaphase, exhibiting transient residency at multiple chromosomes. Chromosomes with inner centromere-localized Plk1 have end-on attached microtubules, diminished Spindle Assembly Checkpoint (SAC) components, and low Shugoshin 1 (Sgo1) levels at the inner centromere. Mechanistically, recruitment is driven by Cdk1 activity and requires Plk1’s Polo-Box Domain (PBD). Moreover, inhibition of Bub1 or Protein Phosphatase 2A (PP2A) increases Plk1 recruitment and residency at the inner centromere. Collectively, our data identify a novel pathway for Plk1 recruitment to the inner centromere that is dynamically regulated by counteracting activities of Cdk1 and Bub1/PP2A.

## Introduction

During mitosis, duplicated sister chromatids, bound by the cohesin complex and attached to microtubules emanating from opposite spindle poles, are accurately segregated into two daughter cells. Segregation errors frequently generate chromosomal instability and aneuploidy, both hallmarks of cancer^1–4^. Therefore, faithful chromosome segregation is carefully orchestrated by molecular signaling at multiple cellular structures including the centrosomes, microtubules, and kinetochores. The kinetochore, a multi-protein complex assembled at the centromere, serves as a structural and signaling hub for various events in mitosis. CENP-A, an alternate Histone 3 (H3), epigenetically specifies centromeres and maintains the location of kinetochore assembly^5^. In early mitosis, the KMN network composed of the Knl1, Mis12, and the Ndc80 complexes is recruited to the Constitutive Centromere Associated Network (CCAN) to form the outer kinetochore^6–8^. The outer kinetochore serves as the scaffold for microtubule binding and chromosome congression to the metaphase plate. Once all chromosomes are correctly attached to microtubules and bioriented, the Spindle Assembly Checkpoint (SAC), recruited to Knl1 is silenced, leading to the activation of the Anaphase Promoting Complex/Cyclosome (APC/C), a multi-subunit E3 ligase^9–11^.

These mitotic signaling events along the centromere and kinetochore are catalyzed by kinases including Cdk1^12^, Bub1^13–16^, Aurora B^12^, Mps1^17^ and Plk1^18–21^. While Mps1 (outer-kinetochore) and Bub1 (centromere) functions are limited to a cellular structure, region of the kinetochore or timing in mitosis, Plk1 performs a variety of functions throughout mitosis across different cellular structures. Prior to mitotic entry, Plk1, itself activated by Aurora A and its co-factor Bora, activates Cdk1^22,23^. Activation of Cdk1-Cyclin B1 complex is essential for mitotic entry. During early mitosis, Plk1 localizes to the chromosome arms and kinetochores, where its activity promotes removal of chromosomal arm cohesion^24–27^ and establishment of kinetochore-microtubule attachments^28,29^, respectively. Additionally, Plk1 cooperates with Mps1 to maintain the SAC, preventing premature mitotic exit^30^. Once the chromosomes are under tension by microtubule pulling forces, Plk1 prevents centromere dislocation and kinetochore disruption^18,20^. To localize to these diverse subcellular structures, Plk1 binds phosphorylated proteins via its Polo-Box Domain (PBD). In early mitosis, Plk1 transduces Cdk1 driven signals through its PBD^31^, increasing the local activity of Plk1 near a substrate leading to a high degree of phosphorylation. The PBD is important for Plk1 recruitment to the kinetochore ^32,33^. As Cdk1 activity is stable in early mitosis, Plk1 function is relatively uniform and persistent, including at the kinetochores. During this stage, the predominant kinetochore Plk1 recruitment is by binding to Cdk1 or Plk1 phosphorylation on CENP-U and Bub1^34,35^ at the inner kinetochore and outer-kinetochore, respectively. Recruitment to these regions is important for microtubule binding and maturation.

In addition to CENP-U and Bub1, Plk1 binds to and/or phosphorylates other proteins found at the centromere (Plk1 Interacting Checkpoint Helicase/PICH)^18,20,36^, inner kinetochore (CENP-Q)^35^, and outer kinetochore (CLASP2)^37^. Since the kinetochore spans ∼100 nm and the Plk1 kinase domain is only ∼5 nm, primary recruitment to CENP-U and Bub1 alone is insufficient to explain Plk1 functions at other regions of the kinetochore. Previously, we proposed a model in which Plk1 is recruited to multiple regions on the kinetochore, generating spheres of activity around each^21^. We also found that a majority of Plk1 was recruited to the centromere region ∼13.9 nm interior to CENP-A during metaphase^21^. Dynamic Plk1 recruitment could explain the presence of these multiple kinetochore interactors, as Plk1 can be rapidly retargeted to distinct locations as mitosis progresses. For example, Plk1 targets the spindle midzone at anaphase onset, regulating cytokinetic furrow ingression^38–40^. However, direct visualization of dynamic retargeting of Plk1 along the length of the early mitotic kinetochore has remained elusive. Although dynamic retargeting is not reported, there are known chances of kinetochore levels of Plk1 which peak at prometaphase and then decline^41^. Moreover, Plk1 kinetochore levels are higher on misaligned chromosomes compared to those correctly aligned at the spindle equator ^42^. Finally, Plk1 creates the 3F3/2 phosphoepitope on a subset of kinetochores—preferentially found on misaligned chromosomes—in a SAC dependent manner^42^. These observations hint at more dynamic recruitment through Plk1, which could result from either positive feedback of Plk1 self-priming, or from negative feedback from phosphatases shifting the Cdk1 phosphorylation equilibrium at PBD binding sites.

In this study, we combine fixed and live imaging with phosphorylation manipulation, proximity labeling, and protein depletion to reveal a conserved, dynamic recruitment of Plk1 to the inner centromere of a subset of chromosomes during prometaphase and metaphase. This recruitment succeeds end-on microtubule attachment, SAC silencing and removal of Shugoshin 1 (Sgo1) from the inner centromeres. We also find that Sgo1-PP2A recruited through Bub1 phosphorylation on Threonine 120 of H2A (H2A T120ph)^43,44^ controls Plk1 recruitment to INCENP in a Cdk1- and PBD-dependent manner. This dynamic recruitment illustrates that Plk1 operates in a distinct inner centromere environment separate from other spheres of activity at the kinetochore.

## Results and discussion

### Plk1 is dynamically recruited to the inner centromere in late prometaphase and metaphase

It is well established that Plk1 is recruited to the mitotic kinetochore, via its PBD, binding Cdk1- and Plk1-generated phosphorylation sites on CENP-U (inner-kinetochore), Bub1 and BubR1 (outer-kinetochore)^45–47^. Through careful microscopic analysis, we identified an addtional focus of Plk1 at the inner centromere of mitotic chromosomes in retinal pigmental epithelial (hTERT-RPE1) cells **(Figure 1A)**. To track the timing of Plk1 recruitment to the inner centromere, we fixed and imaged pre-anaphase mitotic cells. We categorized the cells into prophase, early (early PM) or late (late PM) prometaphase, and metaphase, based on centrosome separation and chromosome alignment along the cell equator. Plk1 inner centromere localization was observed only during late prometaphase and metaphase **(Supplementary Figure 1A**). These findings are not unique to this cell type—we visualized Plk1 inner centromere recruitment during late prometaphase and metaphase in both HeLa (cervical cancer) and MCF10A (untransformed breast) cell lines **(Figure 1A)**, suggesting a generalized mitotic phenomenon. We differentiated the inner centromere Plk1 signal from kinetochore Plk1 on an adjacent chromosome by linescan analysis in single plane images of kinetochore pairs marked with CENP-A **(Figure 1B)**. Interestingly, Plk1 inner centromere recruitment was present in a subset of chromosomes across the three cell lines. Hence, we categorized the chromosomes as Plk1 Inner Centromere positive or negative (Plk1 IC +ve vs Plk1 IC -ve) **(Figure 1C)**. In all, metaphase cells had a higher number of Plk1 IC +ve chromosomes than late prometaphase in all three lines **(Figure 1D)**, suggesting a conservation of this recruitment. Collectively, these data support *bona fide* Plk1 inner centromere recruitment to chromosomes during late prometaphase and metaphase.

**Figure 1.**
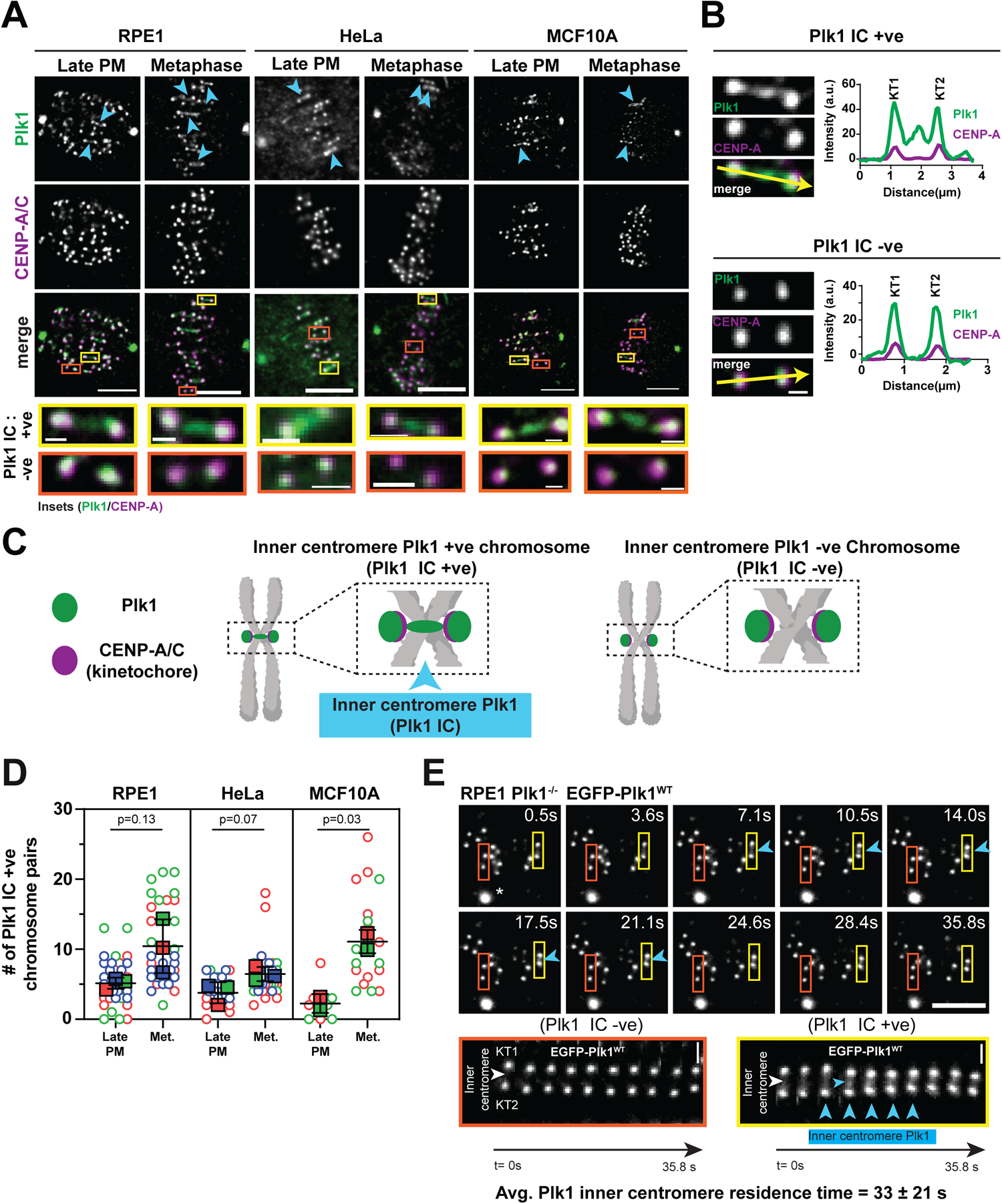
Plk1 is dynamically recruited to the inner centromere in late mitosis. **A)** Representative 5-plane immunofluorescence projection images of Plk1 recruitment to the kinetochore and inner centromere during late prometaphase (late PM) and metaphase in RPE1, HeLa, and MCF10A cells. Kinetochores are marked with CENP-A (RPE1, HeLa) and CENP-C (MCF10A). Arrows indicate Plk1 IC +ve kinetochores. Insets highlight Plk1 IC +ve (yellow) and Plk1 IC -ve (orange) kinetochores. Scale bar-5 µm, Insets-500 nm (RPE1, MCF10A), 1 µm (HeLa). **B)** Single plane linescan analysis of Plk1 and CENPC in a Plk1 IC +ve and Plk1 IC -ve kinetochore from an RPE-1 cell. Scale bar-500 nm. Arrow indicates direction of linescan. **C)** Illustrated chromosome pair highlighting Plk1 kinetochore recruitment and presence (Plk1 IC +ve) or absence (Plk1 IC -ve) at the inner centromere. **D)** Graph depicting average (± SD) number chromosome pairs/cell with inner centromere localized Plk1 from (A). Circles and squares represent individual cells (4-12/replicate) and average values, respectively, from color-coded biological replicates (n= 2). **E)** Individual time frames from live cell imaging of metaphase RPE1 Plk1^-/-^ EGFP-Plk1^WT^ cell. Orange and yellow boxes/insets track kinetochore pairs oscillating along the spindle axis. Blue Arrowheads indicate the presence of inner centromere Plk1. Average values of 12 kinetochore pairs from 5 cells. Data is also graphed in Figure 3I. Time indicated in seconds. Scale bar-5 µm (whole cell), 1 µm (insets).

Given Plk1 is visible at a subset of inner centromeres in fixed cells, we considered this could either be stable recruitment to a few specific chromosomes or ongoing dynamic recruitment to different chromosomes. To assess this, we performed live cell, confocal imaging of RPE1 cells, stably expressing EGFP-tagged wildtype Plk1 (RPE1 Plk1^-/-^ EGFP-Plk1^WT^). As expected, EGFP-tagged Plk1 localized to both kinetochores and centrosomes. Tracking the dynamics of inner centromere Plk1 recruitment at metaphase, we observed dynamic inner-centromere recruitment of Plk1 with an average residing time of ∼33 seconds **(Figure 1E, SuppVideo 1)**. These findings suggest that inner centromere Plk1 recruitment is an ongoing dynamic process intermittently reaching many, if not all, centromeres.

### Dynamic inner centromere recruitment requires the PBD and Cdk1 activity

Plk1 kinetochore recruitment frequently requires PBD binding to Cdk1-phosphorylated motifs on the interacting proteins^33,45^. Point mutations (H538A, K540A) within the PBD abolish this recruitment^31,33,48,49^. To test if inner centromere Plk1 recruitment is also dependent on the PBD, we imaged RPE1 cells simultaneously expressing a EGFP-Plk1 PBD WT and a Flag-Plk1 PBD mutant **(Supplementary Figure 2A)**. Flag-Plk1 PBD mutant remained delocalized even as EGFP-Plk1 PBD WT, as expected, was dynamically recruited to the inner centromere **(Supplementary Figure 2B)**, demonstrating the requirement for a functional phosphopeptide binding. To test if Cdk1 phosphorylation is required for Plk1 inner centromere localization, we briefly inhibited Cdk1 chemically, allowing for mitotic entry but preventing early mitotic exit **(Supplementary Figure 2C)**. In addition to decreasing Plk1 kinetochore recruitment, as previously described^45^, Cdk1 inhibition abrogated inner centromere recruitment **(Supplementary Figure 2D-E)**. These findings demonstrate that Cdk1 activity is important for Plk1 inner centromere recruitment. Given the dynamic recruitment, the requirements for PBD and Cdk1, our findings suggest that the PBD docking site at the inner centromere is in dynamic equilibrium between phosphorylated and non-phosphorylated states.

### Chromosomes with dynamic inner centromere Plk1 have mature spindle attachment

To understand the dynamic nature of Plk1 inner centromere recruitment, we considered the molecular events underpinning prometaphase to metaphase transition. Correction of erroneous kinetochore-microtubule binding to generate end-on attachments is a major molecular event prior to metaphase^11,30,50^, which should coincide with maximal inner centromere Plk1 recruitment. Additionally, Spindle Assembly Checkpoint (SAC) proteins are recruited to unattached kinetochores in early prometaphase, delaying anaphase onset by inhibiting the Anaphase Promoting Complex/Cyclosome (APC/C)^51^. Hence, we decided to observe the status of microtubule binding and SAC on Plk1 IC +ve chromosomes.

First, we probed Astrin, a kinetochore and microtubule binding protein recruited to the outer-kinetochore of chromosomes with end-on microtubule attachment. Dynamically recruited PP1 on Astrin stabilizes the end-on attachment prior to bi-orientation^52^. Plk1 IC +ve chromosomes in late prometaphase and metaphase exhibited 94 ± 5% Astrin recruitment to kinetochores of both sister chromatids **(Figure 2A-B)**.

**Figure 2.**
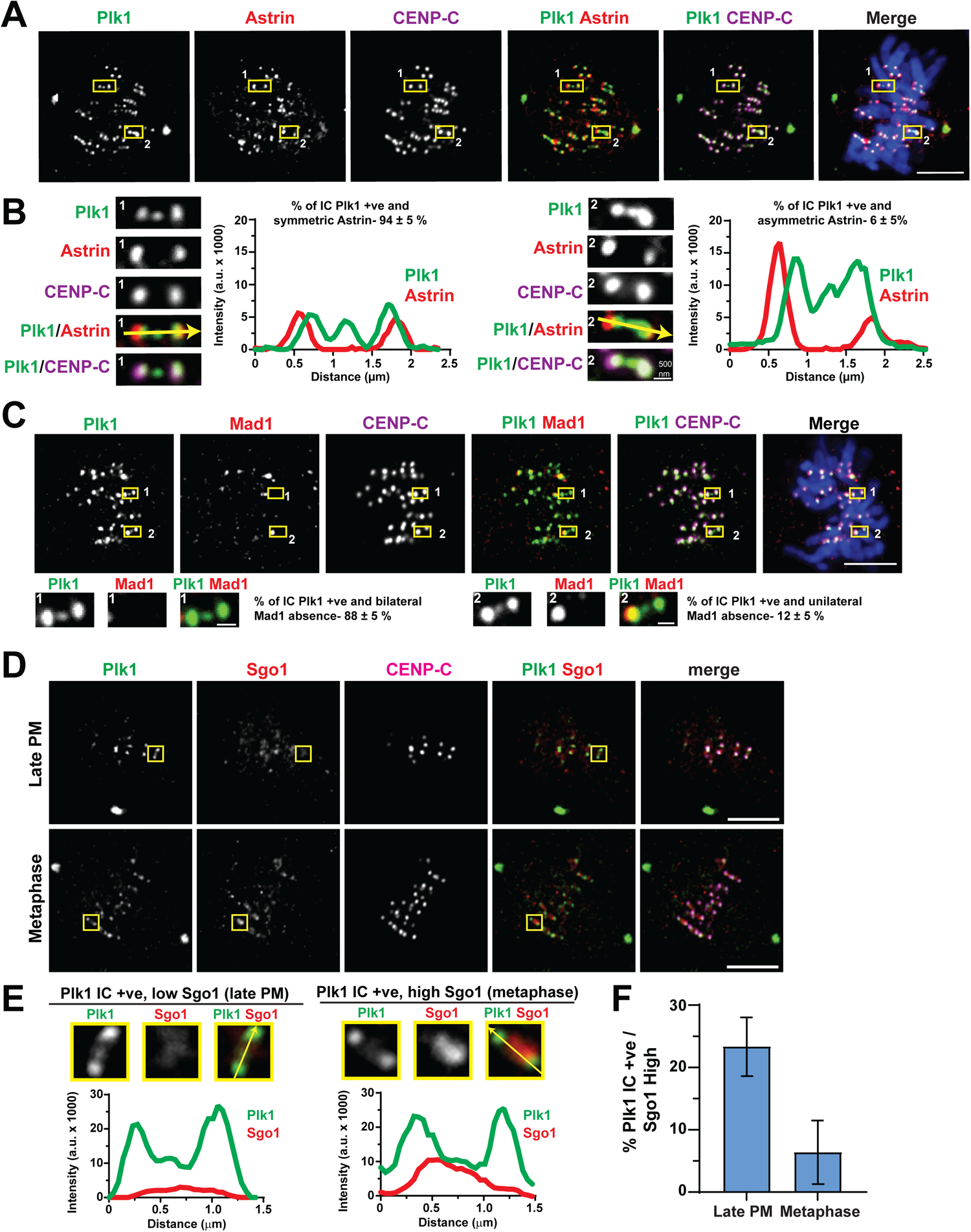
Chromosomes with dynamic inner centromere Plk1 recruitment are bound to microtubules in an end-on manner, lack Mad1, and have reduced Shugoshin 1 (SGO1) **A)** Representative immunofluorescence image and linescan analysis of an RPE1 cell in late PM with examples of symmetric (box1) or asymmetric (box2) Astrin recruitment to Plk1 IC +ve chromosome pairs, denoted by CENP-C. Scale bar-5µm (whole cell), 500 nm (insets). **B)** Linescan analysis and insets from (A). Values indicate average (± SD) Astrin distribution from Plk1 IC +ve chromosomes (n= 280, 2 replicates). Arrow indicates direction of linescan. **C)** Representative immunofluorescence image of an RPE1 cell in late PM with examples of bilateral (box1) or unilateral (box2) Mad1 kinetochore absence on Plk1 IC +ve chromosome pairs, denoted by CENP-C. Values indicate average (± SD) Mad1 distribution from Plk1 IC +ve chromosomes (n= 173, 3 replicates). Scale bar-5 µm (whole cell), 500 nm (insets). **D)** Representative single plane immunofluorescence images of RPE1 cells in late PM and metaphase. Boxes indicate Plk1 IC +ve chromosome pairs, indicated by CENP-C. Scale bar-5µm. **E)** Insets and linescan analysis of chromosome pairs from (C), demonstrating low (< 5000 a.u.) and high (> 5000 a.u.) Sgo1 levels. Arrow indicates direction of linescan. **F)** Graph depicting average (± SD) percentage Plk1 IC +ve chromosome pairs with high (> 5000 a.u.) Sgo1 (n= 20 late PM / 79 metaphase, 2 replicates) of each example

To determine the SAC status of Plk1 IC +ve chromosomes, we labeled Mad1, a component of the SAC found on unattached kinetochores. Mad1 was removed from both kinetochores in 88 ± 5% of Plk1 IC +ve chromosomes **(Figure 2C)**, suggesting SAC silencing and consistent with Astrin recruitment **(Figure 2A-B**).

Since inner centromere Plk1 recruitment succeeds end-on microtubule attachment and SAC silencing, we were curious if distribution of any other inner centromere protein differs between unattached vs. end-on attached chromosomes. Previous studies demonstrate that Sgo1 is redistributed away from the inner centromere in a microtubule-binding and tension-dependent manner^53–55^. Therefore, we tested if inner centromere Plk1 recruitment correlates with Sgo1 redistribution. We observe high Sgo1 levels at almost every inner centromere during prophase **(Supplementary Figure 3A-B)**, however levels varied as mitosis progressed **(Figure 2D-E, Supplementary Figure 3C)**. In late prometaphase, only 23 ± 5% of Plk1 IC +ve chromosomes exhibited high Sgo1, dropping to 6 ± 5% by metaphase **(Figure 2F)**. This illustrates that Sgo1 redistributes away from the inner centromere in chromosomes when Plk1 is recruited. Taken together our data illustrate that inner centromere Plk1 recruitment succeeds end-on microtubule binding, SAC silencing, and inner centromere Sgo1 removal.

### Bub1 and PP2A activity control Plk1 recruitment to the inner centromere

Since dynamic inner centromere Plk1 recruitment occurs through Cdk1-driven PBD binding, we reasoned that the Plk1 interactor transitions between a phosphorylated and a non-phosphorylated state through dynamic kinase and/or phosphatase activity. Indeed, PP2A is recruited through Sgo1, which in turn is recruited by Bub1 phosphorylation of Threonine 120 on H2A (H2A T120ph)^43,44,53,55^. Thus, we hypothesized that Sgo1-recruited PP2A dephosphorylates the inner centromere Plk1 interactor(s) to prevent Plk1 binding prior to metaphase **(Figure 3A)**. If so, Bub1 inhibition would prevent inner centromere localization of Sgo1-PP2A, thereby increasing inner centromere Plk1. To test this, we chemically inhibited Bub1 using BAY-1816032^56^ in cells released from a STLC prometaphase arrest **(Figure 3B)**. We confirmed Bub1 inhibition by loss of H2A T120ph **(Figure 3C)**. Strikingly, Bub1 inhibition doubled the number of Plk1 IC +ve metaphase chromosomes (19.3 ± 1%) compared to untreated controls (8.8 ± 1.4%) **(Figure 3D)**. Furthermore, PP2A inhibition using Okadaic Acid (OA) also significantly increased the number of Plk1 IC +ve metaphase chromosomes **(Figure 3E-F)**. Combined, our results suggest a model where Sgo1 removes inner centromere Plk1 in a PP2A-dependent manner.

**Figure 3.**
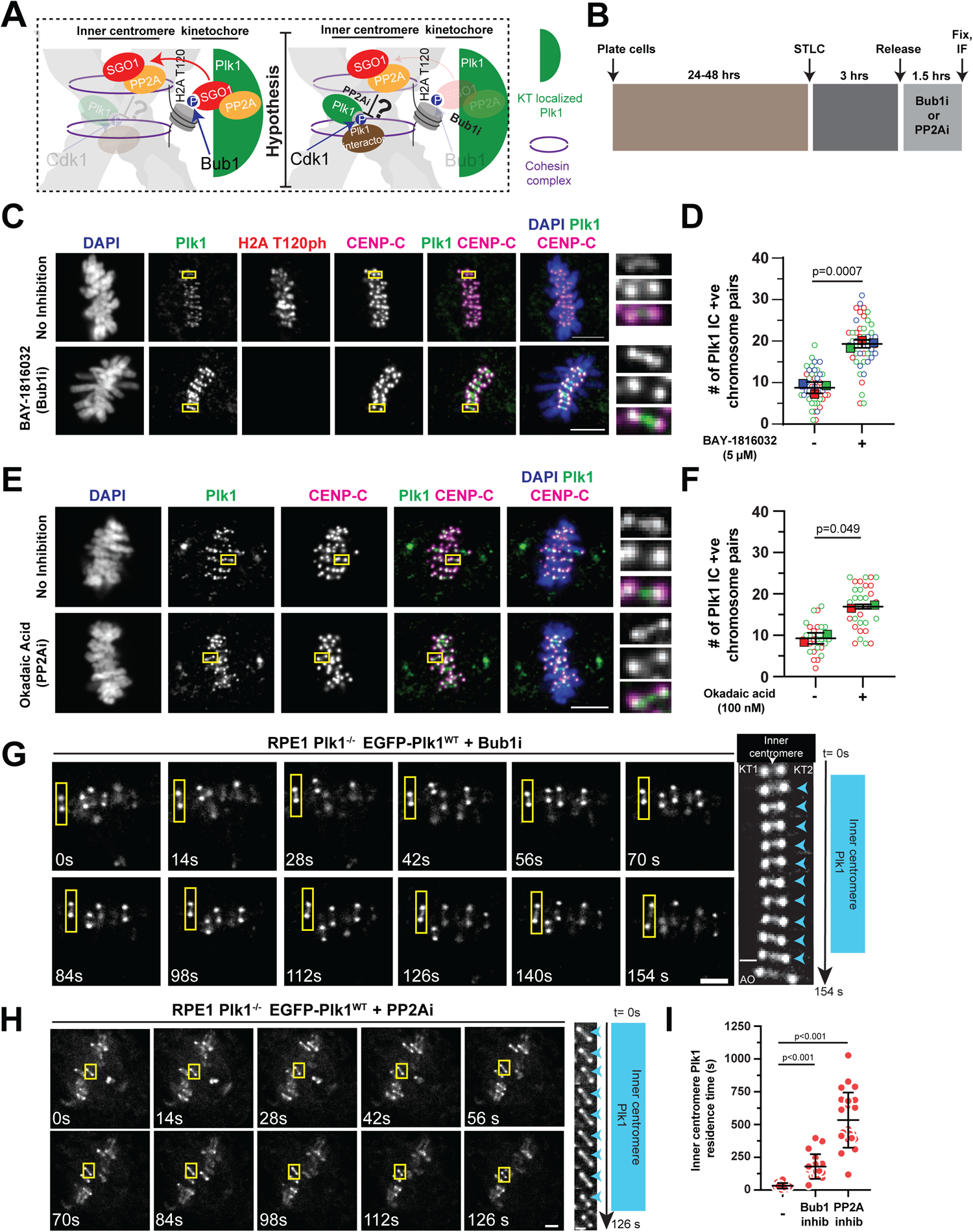
Bub1 and PP2A activity control Plk1 recruitment to the inner centromere. **A)** Schematic illustrating SGO1-PP2A recruitment to kinetochore through Bub1 phosphorylation of H2A T120 and hypothesized model that activity counters Cdk1 recruitment of Plk1 to inner centromere. **B)** Experimental setup for panels C-F. **C)** Representative immunofluorescence images of metaphase RPE1 cells with or without Bub1 inhibition. Insets highlight Plk1 IC +ve chromosome pair. Scale bar-5µm. **D)** Graph depicting average (± SD) number chromosome pairs/cell with inner centromere localized Plk1. Circles and squares represent individual chromosome pairs (15/replicate) and average values, respectively, from color-coded biological replicates (n= 3). **E)** Representative immunofluorescence images of metaphase RPE1 cells with or without Okadaic Acid treatment. Insets highlight Plk1 IC +ve chromosome pair. Scale bar-5 µm. **F)** Graph depicting average (± SD) number chromosome pairs/cell with inner centromere localized Plk1. Circles and squares represent individual chromosome pairs (14/replicate) and average values, respectively, from color-coded biological replicates (n= 2). **G-H)** Individual time frames from live cell imaging of metaphase RPE1 Plk1^-/-^ EGFP-Plk1^WT^ cells with Bub1 (**G**) or PP2A (**H**) inhibition. Yellow boxes/insets track kinetochore pairs oscillating along the spindle axis. Blue Arrowheads indicate the presence of inner centromere Plk1. Scale bar-2.5 µm. Time indicated in seconds. **I)** Graph depicting average (± SD) residence time of inner centromere Plk1 (18-20 pairs from 3-5 cells) from (D-E). Untreated cell data from Figure 1E.

To understand how Bub1 or PP2A inhibition affect the dynamics of inner centromere Plk1 recruitment, we examined residence time in RPE1 Plk1^-/-^ EGFP-Plk1^WT^ cells under Bub1 or PP2A inhibition. Both Bub1 (180 ± 94 s) and PP2A (534 ± 210 s) inhibition increased inner centromere Plk1 residence time compared to untreated controls (33 ± 21 s) **(Figure 3G-I**, **Figure 1E, SuppVideo 1-3)**. This suggests that increases in Plk1 IC +ve chromosomes after Bub1 or PP2A inhibition result from a decreased removal rate once recruited. Collectively, these data suggest that the inner centromere recruitment of Plk1 is controlled by Bub1 and PP2A activity at the inner centromere.

### INCENP recruits Plk1 to the inner centromere

The PBD-dependent recruitment of Plk1 to the inner centromere suggests phosphorylation-mediated binding to protein(s) in the region. To identify proteins proximal to Plk1 at the inner centromere, we generated cell lines that fused TurboID, a promiscuous biotin ligase with a biotinylation radius of ∼15 nm^57^, to Plk1 with an intact (Plk1-PBD WT) or mutated (Plk1-PBD Mutant) PBD **(Figure 4A)**. As expected Plk1-PBD WT biotinylated both kinetochore and centrosome proteins, whereas the Plk1-PBD mutant only biotinylated centrosome proteins, albeit at a lower level **(Figure 4B)**.

**Figure 4.**
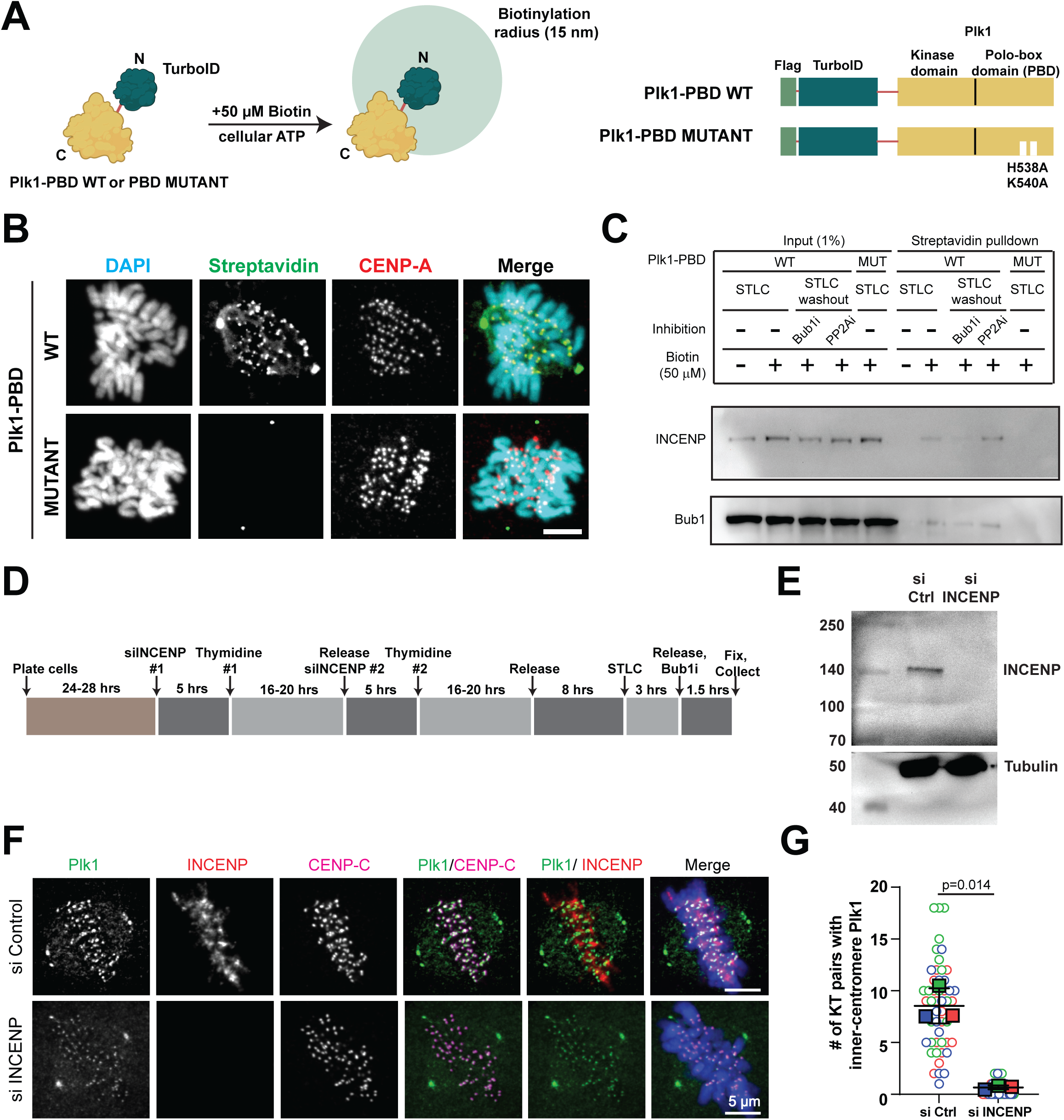
INCENP recruits Plk1 to the inner centromere. **A)** Schematic of proximal labeling with biotin. Domain architecture of Flag-TurboID tagged Plk1 with WT (PBD WT) or mutated (PBD MUTANT) Polo-Box Domain. **B)** Representative immunofluorescence image of RPE1 cell lines stably expressing Flag-TurboID-Plk1 PBD WT or PBD MUTANT and incubated in 50 μM biotin. Alexa Flour 488-conjugated Streptavidin detects proteins proximal to Plk1. CENP-A indicates kinetochore. Scale bar-5 µm. **C)** Immunoblot of protein extracts post-streptavidin pulldown from RPE1 cells expressing Flag-TurboID-Plk1 PBD WT or PBD MUTANT and incubated in 50 μM biotin. Cells were arrested in prometaphase using STLC (STLC) or released from this arrest for progression into metaphase (STLC washout). In STLC washout the Bub1 (5 μM BAY-1816032) or PP2A inhibitor (200 nM OA). Membranes probed for INCENP and Bub1 (known Plk1 interactor) in each condition. **D)** Schematic of the INCENP siRNA experimental setup for E-G. **E)** Immunoblot of protein lysates after non-targeting (Control) or INCENP siRNA depletion. Membranes probed for **I**NCENP and tubulin (loading control). **F)** Representative images of RPE1 cells after siRNA depletion. **G)** Graph depicting average (± SD) number chromosome pairs/cell with inner centromere localized Plk1 from (F). Circles and squares represent individual cells (9-18/replicate) and average values, respectively, from color-coded biological replicates (n= 3).

Previous studies show that INCENP, a component of the CPC, binds Plk1 in a PBD-dependent manner^58^. To test if Plk1 and INCENP co-localize at the inner centromere during metaphase, we used our PBD WT and mutant lines to pull down biotinylated proteins in prometaphase (STLC) or metaphase (STLC washout). To enhance inner centromere Plk1, we inhibited Bub1 or PP2A activity. We identified INCENP in streptavidin pulldown from the PBD WT, but not the PBD mutant line (**Figure 4C**), indicating that Plk1 localizes proximal to INCENP in a PBD-dependent manner. PP2A inhibition increased the amount of biotinylated INCENP **(Figure 4C)**, consistent with the increase in inner centromere Plk1 frequency **(Figure 3C)**. To test if INCENP recruits Plk1 to the inner centromere, we depleted INCENP in RPE1 cells using siRNA. Upon INCENP depletion, recruitment of Plk1 to the inner centromere was abolished **(Figure 4D-F)**. Hence, we conclude that INCENP recruits Plk1 to the inner centromere during metaphase.

### Dynamic Plk1 recruitment to the inner centromere is mediated by Cdk1, Bub1 and PP2A activity

Here, we describe dynamic Plk1 recruitment to the inner centromere, distinct from its kinetochore recruitment as evidenced by microscopic analysis. Curiously, this inner centromere recruitment begins during late prometaphase in a subset of chromosomes, briefly lasting until late metaphase (**Figure 1**). Localization requires end-on microtubule attachments, SAC silencing, and Sgo1 inner centromere removal **(Figure 2)**. Furthermore, Cdk1 activity promotes recruitment, whereas Bub1-mediated Sgo1-PP2A activity opposes it **(Figure 3)**. Finally, we find INCENP to be necessary for Plk1 inner centromere localization **(Figure 4)**.

Plk1 is recruited to every kinetochore during mitosis, primarily to Bub1 and CENP-U^29,45,47^, sites that permit it to phosphorylate multiple substrates. Specifically, it acts to stabilize KT-microtubule attachments in a SAC-dependent manner^29,59^. In contrast to this stable recruitment, we find Plk1 is dynamically recruited to the inner centromere, typically when kinetochores are attached in an end-on manner and the SAC has been silenced.

Why do only certain chromosomes have Plk1 recruitment to the inner centromere? One possibility is that Plk1 is recruited to the inner centromere of all chromosomes, but our imaging conditions lack the sensitivity and sampling frequency to detect Plk1. Perturbations with Bub1 inhibition and Okadaic acid support this assertion, as the recruitment becomes visible in all. A second answer is that only some chromosomes require active Plk1 at the inner centromere. Indeed, recent studies have found that larger and more peripheral chromosomes are more likely to be missegregated or to delay mitosis^60–62^. Additionally, kinetochore size correlates with chromosome missegregation frequencies^63^. Thus, additional mechanisms, which could require Plk1 recruitment, could fire at specific kinetochores to assure high fidelity of chromosome segregation.

INCENP is necessary for Plk1 inner centromere recruitment. In Drosophila, Polo and INCENP co-localize with INCENP important for Polo phosphorylation at Thr 182^64^. In mammals, the importance of Plk1 Interaction between INCENP and the Plk1-PBD through Thr 412 was shown previously *in vitro*^58^. A T412A mutation abolished inner centromere recruitment and modestly delayed anaphase onset^58^. Previous biochemical studies could not illustrate the temporal nature of this or other inner centromere phosphorylations during mitotic progression. Our study adds that this site, or others, dynamically recruit Plk1 to the inner centromere at mature KTs.

We report INCENP-dependent dynamic docking of Plk1 at the inner centromere, a process that has not been directly observed in live cells likely due to low levels, and a fraction of chromosomes. The dynamic recruitment is apparently controlled via a Cdk1-dependent docking site, and is also controlled by Bub1 and PP2A activities. It will be important to determine whether phosphorylated T412 is solely responsible for recruitment, and which specific mitotic errors arise due to inability to recruit Plk1, or inability to dynamically alter levels.

## Materials and methods

### Cell line derivation and culture procedures

#### Cell culture

All cells were maintained at 37°C and 5% CO2 in a humidified, water-jacketed incubator and propagated in media supplemented with 10% fetal bovine serum (GeminiBio, 900-108), 100 units/ml penicillin-streptomycin (Gibco, 15070063), and 5 ug/ml plasmocin prophylactic (Invivogen, ant-mpp). hTERT-RPE1 and derived cells were cultured in a 1:1 mixture of Dulbecco’s Modified Eagle’s Medium (DMEM) and Ham’s F-12 media supplemented with 2.5 mM L-glutamine (Cytiva Hyclone, SH3026101). HeLa and Phoenix cells were cultured in DMEM supplemented with 4.0 mM L-glutamine and 4,500 mg/l glucose (Cytiva Hyclone, SH3024301). MCF10A cells were cultured in “bullet kit” media: DMEM/F-12 media, 5% horse serum (Gibco, 16050122), 20 ng/ml EGF (Peprotech, AF-100-15), 0.5 mg/ml hydrocortisone (MP Biomedicals, 0219456901), 100 ng/ml cholera toxin (Enzo Life Sciences, BMLG1170002), and 10 ug/ml insulin (Millipore Sigma, I9278). All cell lines were tested for mycoplasma contamination with the MycoAlert Mycoplasma Detection Kit (Lonza, LT07-418).

#### Plasmid construction

TurboID sequence was amplified from a pcDNA3 plasmid containing 3xHA-TurboID-NLS (Addgene #107171). Flag tag sequence was added to the N-terminus using a forward primer. This amplicon was inserted into a pQC backbone between NotI and AgeI sites upstream of Internal Ribosome Entry Site (IRES) under a CMV promoter. This plasmid backbone contains a Neomycin resistance marker downstream of IRES. Plk1 WT or Plk1 PBD mutant (H538A and K540A) sequence was cloned into this backbone on the C-terminus of TurboID between AgeI and EcoRI sites to generate the fusion construct.

#### Retroviral transgenesis

For stable retroviral transduction, constructs were co-transfected with a VSV-g pantropic envelope vector into Phoenix cells, a HEK293 cell derivative expressing gag and pol genes from Moloney murine leukemia virus, using FuGene® HD (Promega, PR-E2311) transfection reagent. 48 h post-transfection, media containing viral particles was collected, clarified by centrifugation, filtered through a 0.45 μm membrane and diluted 1:1 with complete medium containing 10 ug/ml polybrene (Millipore, TR-1003-G). Target cells (hTERT-RPE1 Plk1^flox/Δ^, containing one floxed and one deleted Plk1 allele^39^) were infected at 30-50% confluence for 24 h and then selected under 0.4 mg/mL Geneticin/G418 sulfate (Gibco, 11811-023) for an additional 5 days. Surviving cells were further purified by limiting dilution to obtain individual clones. Construct expression and localization was validated by immunoblotting and immunofluorescence.

#### siRNA transfection

RPE1 cells were plated in 6-well plates using antibiotic-free media. At 50-60% confluency, cells were transfected with 20 nM siRNA targeting INCENP (IDT, hs.Ri.INCENP.13.1) or negative control siRNA (IDT, 51-01-14-04) using Lipofectamine™ RNAiMAX (Invitrogen, 13778075) according to manufacturer’s instructions. After 5 h, media was replaced with 2.5 mM thymidine. After 16-20 h, media was removed, and cells were transfected a second time with 20 nM siRNA as described above. After 5 h, media was removed, cells were trypsinized and replated on coverslips (for immunofluorescence imaging) or 6-well plates (for immunoblotting). 5 h later, cells were challenged a second time with 2.5 mM thymidine. After 16-20 h, cells were rinsed once with HBSS (Cytiva Hyclone, SH3003102), twice with fresh media, and then released for 8 h in fresh media. Cells were then challenged with 5 μM S-Trityl-L-cysteine (STLC) for 3 h, rinsed once with HBSS, then challenged with 5 μM BAY1816032. After 1.5 h, cells were fixed for immunofluorescence imaging or collected, pelleted, and cryopreserved for immunoblotting.

#### Chemicals

Chemicals and concentrations used in this study: BAY1816032 (SelleckChem, 1816032, 5 μM), Okadaic Acid (Sigma, 09381, 100 or 200 nM), RO-3306 (Tocris, 41811, 10 μM), STLC (Santa Cruz Biotech, sc-202799, 5 μM), Thymidine (Millipore, 6060, 2.5 mM).

### Proximity Labeling and pulldown

To prepare biotin free media, 500 mL of cell culture media supplemented with serum, antibiotic and antimycotic was incubated with 500 µL of Streptavidin agarose beads (Millipore Sigma, #69203-3) and agitated on a circular shaker at 4°C for three days. Post-incubation, the media was vacuum filtered using a 0.2 µm membrane (Millipore Sigma, #S2GPU05RE). Prior to biotinylation, normal culture media was replaced with biotin-free media and the cells were cultured for 48 h. Cells were synchronized in mitosis using 5 µM STLC for 6 h. For washout samples, cells were released into media containing 5 μM BAY1816032 or 200 nM Okadaic acid for 1.5 h. For biotinylation, cells were incubated with media containing 50 µM biotin (Sigma Aldrich, #B4639) for 10 min at 37°C and 5% CO_2_ for 10 min. Cells were then processed for immunofluorescence microscopy (see microscopy section) or pulldown (see below) and Western blotting (see immunoblotting section).

For pulldown, mitotic cells were isolated using shake-off and lysed in a buffer containing 50 mM Tris pH 7.4, 150 mM NaCl, 1% NP-40, 1 mM EGTA, 1.5 mM MgCl_2_, 1mM PMSF and protease inhibitor cocktail (Fisher, #78430). Total protein was quantitated by *DC*™ Protein Assay Kit (BioRad, 5000116) using BSA as standard. 500 µg of total protein lysate was incubated overnight with 50 µL of Streptavidin coated magnetic beads slurry (Fisher, #88817) in 250 µL of lysis buffer (described above) at 4°C on a circular shaker. The beads were washed three times in the lysis buffer. A magnetic rack (Biorad, 1614916) was used to separate the supernatant from the beads. Biotinylated proteins were eluted using 50 µL of Laemmli sample buffer (62.5 mM Tris-HCl pH 8.6, 4% SDS (w/v), 20% glycerol (v/v), 0.01 % bromophenol blue (w/v),10% (v/v) beta-mercaptoethanol) supplemented with 25 mM Biotin at 95°C for 10 minutes.

### Immunoblotting

#### Proximity labeling / Pulldown

Biotinylated proteins were resolved by SDS-PAGE and transferred to 0.2 µm nitrocellulose membrane (Cytiva Amersham™ Protran™, 10600094) at 30V for 16 h in transfer buffer (25 mM Tris, 192 mM Glycine, 20% methanol(v/v), 0.05% SDS (v/v)). The membrane was blocked using 3% BSA in 1x TBST (1x TBS + 0.1% Tween 20) for 30 min at room temperature with gentle agitation. Primary antibodies were diluted in 5% non-fat dry milk in 1x TBST and incubated overnight at 4°C under gentle agitation. The membrane was washed 3x times with 1x TBST for 10 min each under gentle shaking conditions followed by incubation using appropriate secondary antibodies at room temperature for 45 minutes. The membrane was washed and developed with luminol/peroxide (Millipore, WBKLS0500) and visualized with an iBright CL-750 imager (Invitrogen). Image cropping was performed in Illustrator 2024 (Adobe).

#### siRNA

Cell pellets were lysed in buffer (50 mM Tris pH 7.4, 150 mM NaCl, 0.5% sodium dodecyl sulfate, 1% NP-40, 1 mM EGTA, 1.5 mM MgCl_2_) containing 1mM PMSF, 1x HALT™ protease inhibitor cocktail (Thermo Scientific, 1860932) and 1 mM dithiothreitol (Fisher BioReagents™, PR-V3151). Total protein was quantitated by *DC*™ Protein Assay Kit (BioRad, 5000116) and 50 μg separated by SDS-PAGE, transferred to Immobilon®-P PVDF membrane (Millipore, IPVH00010), and blocked for 60 min in 4% nonfat dry milk and 0.1% Tween-20 Tris buffered saline ph 7.4 (TBST+milk). Membranes were incubated with gentle agitation overnight at 4°C with primary antibodies (see Antibodies section) diluted in TBST+milk, washed 3x with TBST, incubated for 1 h at room temperature in secondary antibodies conjugated to horse radish peroxidase diluted 1:10,000 in TBST+milk. Membranes were washed and developed with luminol/peroxide (Millipore, WBKLS0500) and visualized with an iBright CL-750 imager (Invitrogen). Image cropping was performed in Photoshop 2024 (Adobe).

### Microscopy

#### Immunofluorescence imaging and analysis

Cells were plated on 12 mm #1.5 coverslips (Fisher, 1254581P) at 30-50% confluence and cultured for 24-48 h. Unless otherwise specified in the figures, asynchronous cells were used.

Cells were fixed with 3% PFA in PHEM buffer (60 mM PIPES, 27.2 mM HEPES,10 mM EGTA, 8.2 mM MgSO_4_) for 10 mins at 37°C, washed three times in PHEM, permeabilized in PHEM + 0.25% Triton X-100 for 5 min at room temperature (RT), washed three times in PHEM, and blocked for 30 min at RT in 0.1% bovine serum albumin (BSA) and 0.1% Triton X-100 in PBS (PBSTx). Primary antibodies (see Antibodies section) were pooled and diluted in PBSTx + BSA and cells were incubated in for 1 hour at 37°C in a humid chamber to prevent evaporation. Cells were then washed three times in PBSTx. Alexa Fluor (Molecular Probes) secondary antibodies were pooled and diluted at 1:350 in PBSTx + BSA and cells were incubated for another 45 minutes at 37°C in a humid chamber, followed by three washes in PBSTx. Cells were counterstained with DAPI (Sigma Aldrich, 62248) and coverslips were mounted onto glass slides with ProLong™ Diamond anti-fade medium (Fisher, P36970) after drying. Mounted slides were incubated overnight at room temperature before imaging.

Image acquisition was performed on an Eclipse Ti2-E inverted microscope (Nikon) equipped with a CSU-W1 spinning disc confocal scanning unit (Yokogawa), Orca Flash4.0 V2+ digital sCMOS cameras (Hamamatsu), and a high-power laser unit (100 mW for 405, 488, 561, 640 nm wavelength). Optical sections were acquired at 200-300 nm intervals using Elements software (Nikon, version 5.20) and 60x/1.5NA or 100x/1.5NA (both Plan Apo) oil immersion objective. Image analysis and panel cropping was performed in Elements (Nikon, version 4.60). Panels were assembled with overlays using Illustrator 2024 (Adobe).

To determine inner centromere-localized Plk1, optically sectioned cells were visually scanned to capture every kinetochore pair, marked by CENP-C or CENP-A foci. Pairs were considered Plk1 IC +ve if the region between the two kinetochore foci met two criteria. First, Plk1 signal intensity in the region was greater than surrounding background intensity. Second, the region lacked CENP-C or CENP-A signal, which suggested a kinetochore pair from an adjacent plane.

#### Live cell imaging and analysis

Cells were plated in a 35 mm glass bottomed dish (Ibidi, 81156) at 30% confluence and cultured for 48 h before imaging. For untreated condition, asynchronous cells entering mitosis were selected. For Bub1 or PP2A inhibition, cells were incubated with 5 μM BAY1816032 or 200 nM Okadaic acid, respectively, for 2 hours prior to imaging.

Image acquisition was performed on an Eclipse Ti2-E inverted microscope (Nikon) equipped with a CSU-W1 spinning disc confocal scanning unit (Yokogawa), Orca Flash4 digital sCMOS cameras (Hamamatsu), and a high-power laser unit (100 mW for 405, 488, 561, 640 nm wavelength). Environmental control was maintained by a humidified, stage-top chamber (Tokai Hit) set to 37°C and 5% CO_2_. Cells were optically sectioned partially, to avoid photobleaching, at 200 nm intervals every 3-15 s using Elements software (Nikon, version 5.20) and a 60x/1.5NA (Plan Apo) oil immersion objective. Image analysis and panel cropping was performed in Elements (Nikon, version 4.60). Panels were assembled with overlays using Illustrator 2024 (Adobe).

To quantify the residence time, optically sectioned cells were visually scanned to identify kinetochore pairs, marked by EGFP-Plk1. T0 and Tn were marked by inner centromere EGFP-Plk1 intensity that was greater than, and similar to, surrounding background intensity, respectively. Residence time was determined by Tn-T0. Identical contrast values were used for each frame of a given kinetochore pair.

#### Antibodies

The primary antibodies used in this study: Mouse IgG2a anti-Plk1 F8 (Santa Cruz Biotech, sc-17783, 1:250), Mouse IgG1 anti-CENP-A (Enzo life sciences, #ADI-KAM-CC006-E, 1:250), Guinea pig anti-CENP-C (MBL, 101731-882, 1:500), Rabbit anti-Astrin (Atlas antibodies, HPA042027, 1:50), Rabbit anti-Mad1 (gift from Beth Weaver, 1:2000), Mouse IgG2a SGO1 (Abcam, ab58023, 1:250), Rabbit anti-GFP (MBL,598, 1:500), Rabbit anti-H2A T120ph (Active motif, 61195, 1;1000), Mouse IgG1 anti-INCENP (Millipore, 05-940, 1:250-IF 1:1000-WB) and Rat anti-Tubulin (EMD Millipore, MAB1864, 1:10,000-WB). Fluorophore conjugated streptavidin (Streptavidin, Alexa Fluor™ 488 (S32354), 546 conjugate (S11226)) was used to label biotinylated proteins.

### Statistical Analysis

Data analysis was performed using Prism 9 (GraphPad). Unpaired, two-tailed t-test with Welch’s correction was used to compare the means between two populations. P< 0.05 was considered statistically significant.

## Acknowledgements

This work was supported by NIGMS RO1 AAJ1141 (to M.E.B). The authors thank Aussie Suzuki and Beth Weaver for reagents and the Burkard, Weaver and Suzuki labs for helpful suggestions.

---

**Supplementary Figure 1.**
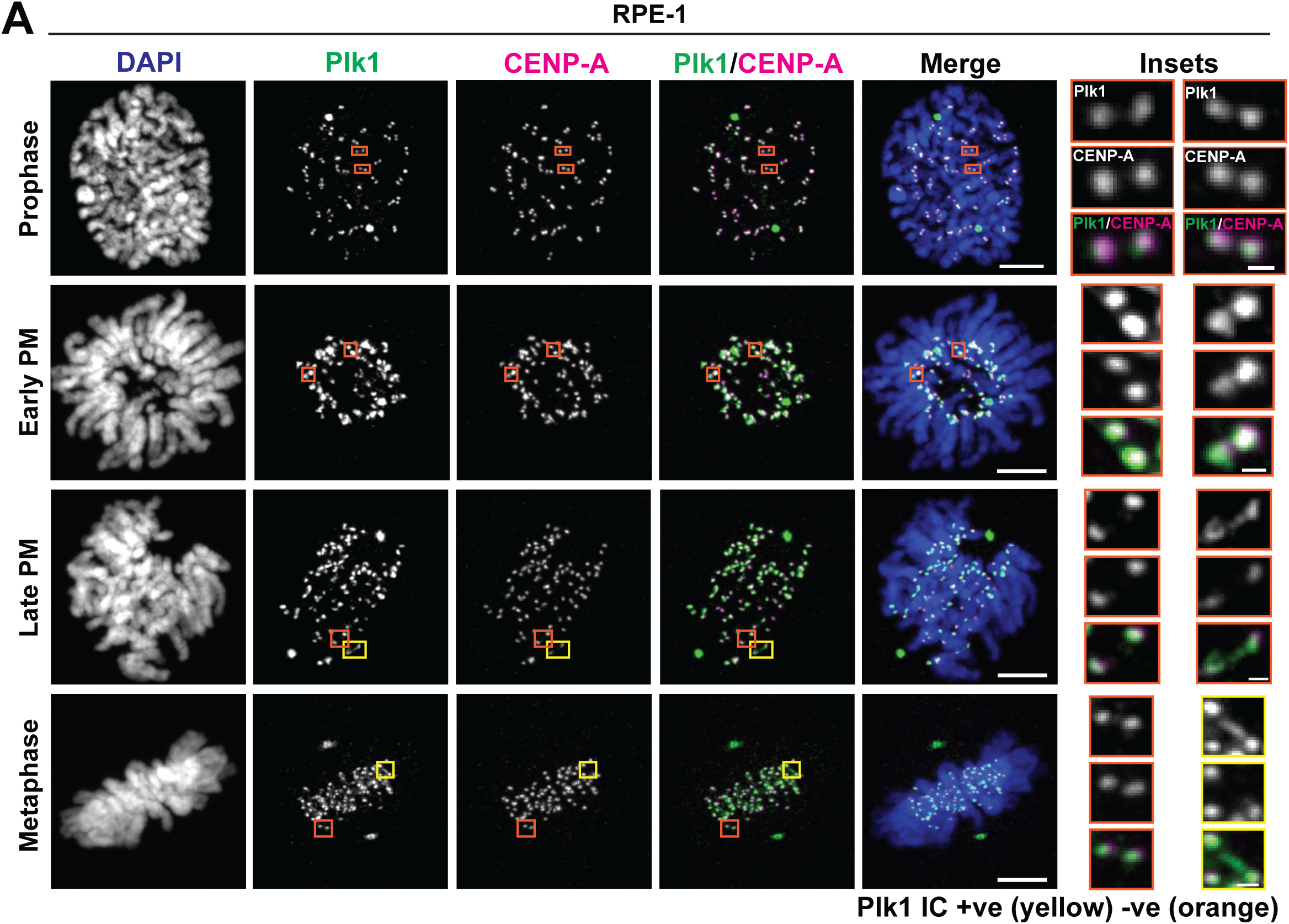
Plk1 localizes to the inner centromere during late prometaphase and metaphase. **A)** Representative immunofluorescence images of Plk1 localization in RPE-1 cells during prophase, early prometaphase (early PM), late prometaphase (late PM) and metaphase. Kinetochore pairs are denoted using CENP-A. Plk1 localization to the inner centromere is observed during late PM and metaphase. Insets bordered with yellow, or orange represent Plk1 IC +ve or Plk1 IC -ve kinetochores, respectively. Scale bar-5µm, Insets-500 nm.

**Supplementary Figure 2.**
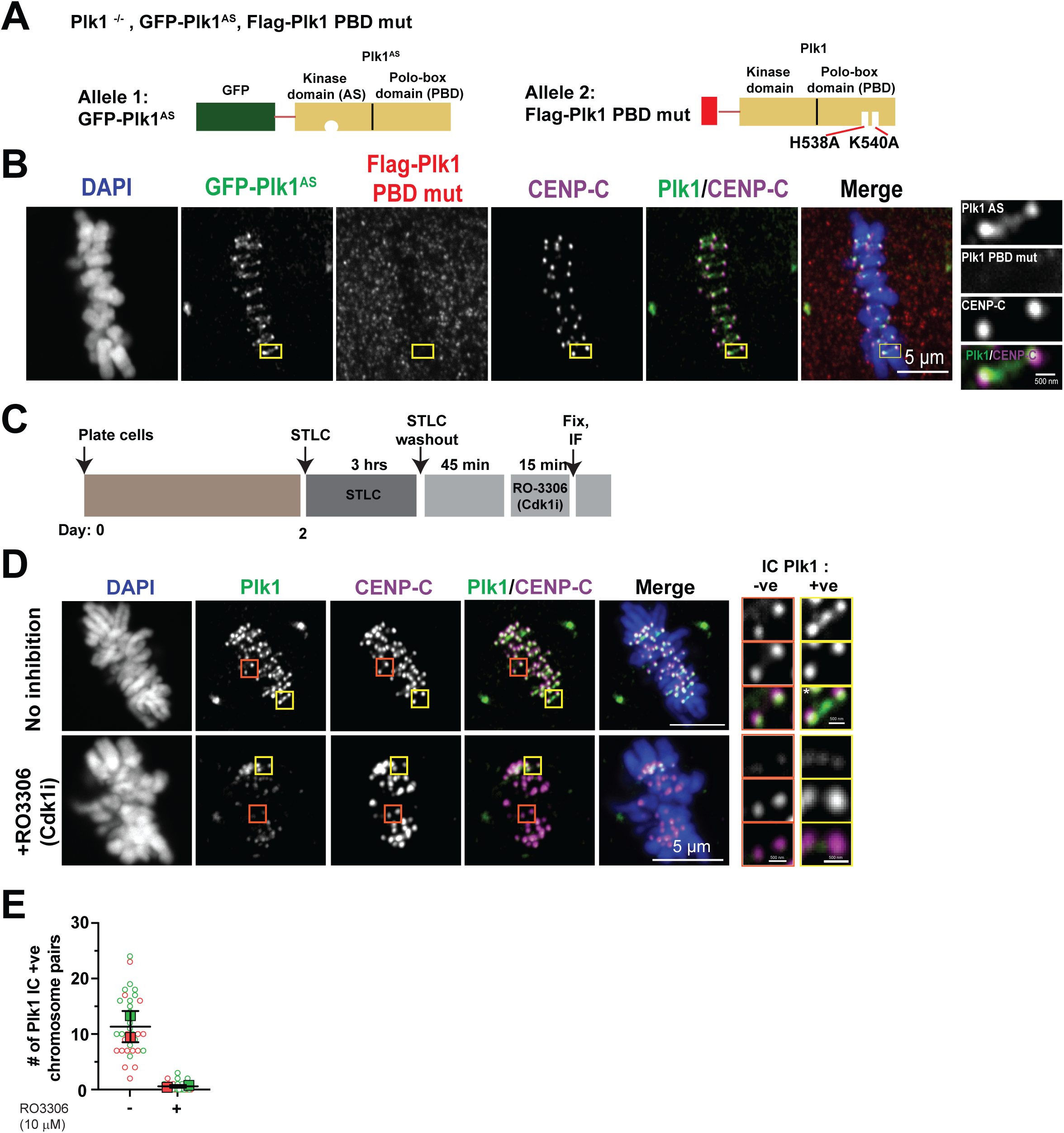
Cdk1 primes the PBD-dependent Plk1 localization to the inner centromere. **A)** Schematic representation of the RPE cell line (Plk1^-/-^ GFP-Plk1^AS^, Flag-Plk1 PBD mut) where the endogenous Plk1 is inactivated while supplementing with alleles of GFP-tagged analogue sensitive Plk1 (GFP-Plk1^AS^) and Flag-tagged Plk1 PBD mutant (H538A, K540A). **B)** Representative immunofluorescence image of RPE1 Plk1^-/-^ GFP-Plk1^AS^, Flag-Plk1 PBD mut cell denoting inner centromere localized GFP-Plk1^AS^ and delocalized Flag-Plk1 PBD mutant. The kinetochore pairs are labeled using CENP-C. Scale bar-5 µm, Insets-500 nm. **C)** Schematic of the experimental setup to study the role of Cdk1 in inner centromere Plk1 recruitment. **D)** Representative immunofluorescence image of Plk1 localization to the inner centromere in RPE-1 cells treated with vehicle (no inhibition) or 10 µM RO-3306 (Cdk1 inhibitor). CENP-C denotes the kinetochore pairs. Scale bars-5 µm, Insets-500 nm. **E)** Graph depicting average (± SD) number chromosome pairs/cell with inner centromere localized Plk1 from (D). Circles and squares represent individual and average values respectively, from color-coded biological replicates (n= 2).

**Supplementary Figure 3.**
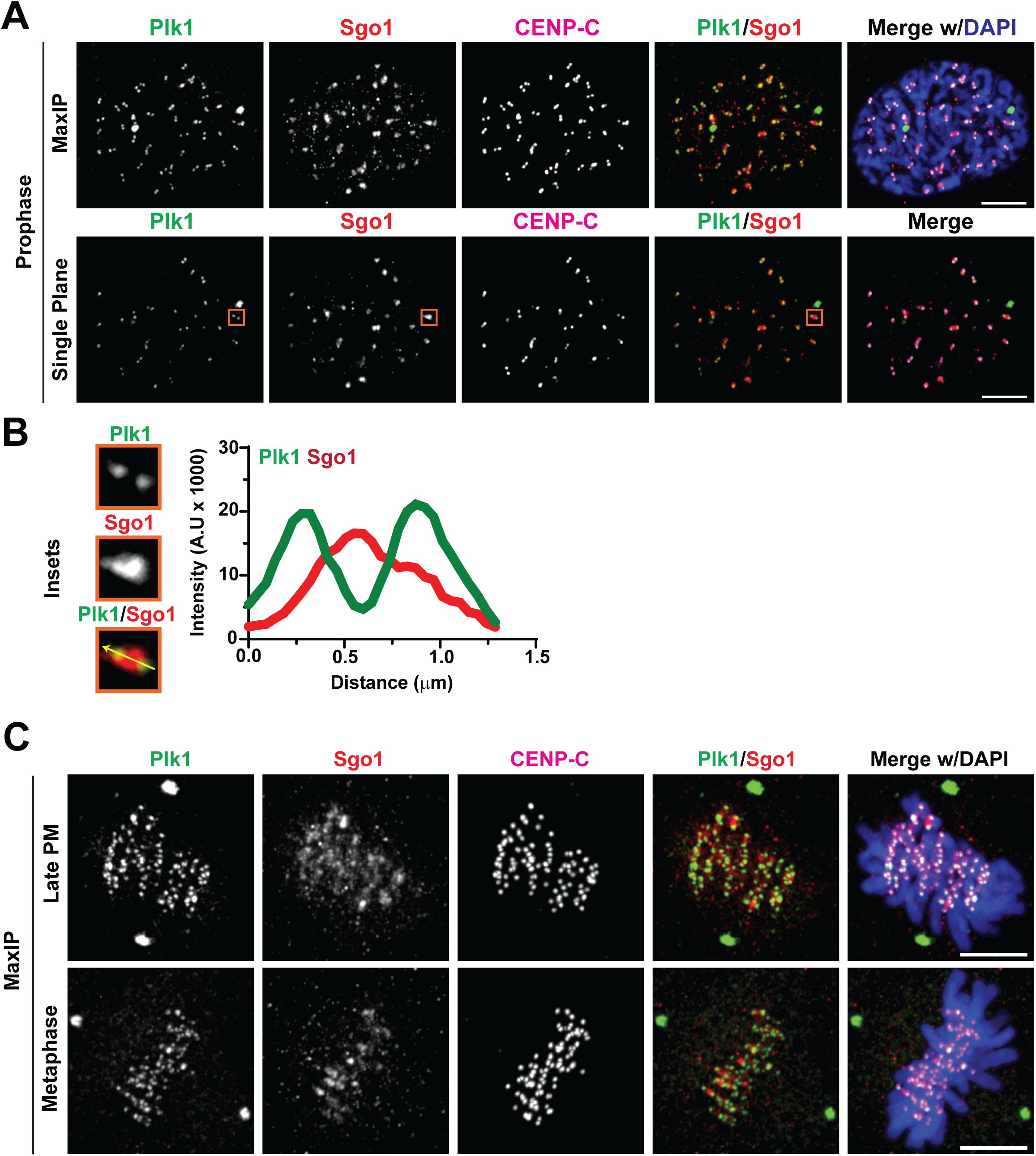
SGO1 level at the inner centromere changes during mitotic progression. **A)** Representative immunofluorescence image of RPE-1 cell denoting Shugoshin-1 (SGO1) localization to the inner centromere in prophase. Single plane and maximum intensity projection (MaxIP) are shown. Kinetochore pairs are marked with CENP-C. Scale bar-5 µm. Orange box represents a kinetochore pair depicting SGO1 and Plk1 localization. **B)** Insets from (**A**) and the linescan depicting Plk1 and SGO1 intensities. Representative MaxIP images of RPE-1 cells in late PM and metaphase depicting SGO1 levels at the inner centromere. Kinetochore pairs are marked using CENP-C. Scale bar-5 µm.

## Notes

### Competing Interest Statement

M.E.B. declares the following: Research funding from Abbvie,
Arcus, Apollomics, Elevation Oncology, Endeavor, Genentech, Puma, and Loxo
Oncology, Seagen. The other authors declare no competing interests.

